# *FMRF* Gene Expression in the Nervous System of the Squid *Doryteuthis Pealei*\* Hatchling

**DOI:** 10.1101/684001

**Authors:** J. Peter H. Burbach, Philip Grant, Stephen Senft, Lizzie Kripke, Anita J.C.G.M. Hellemons, Harish C. Pant

## Abstract

FMRFamide is a neuropeptide that is widely distributed in invertebrates and known to be involved in many physiological functions. Previously we noted marked differences in expression of the *fmrf* gene in the stellate ganglion of *Doryteuthis pealei\** compared to the central nervous system. In this study we aimed to examen the brain systems of *Doryteuthis pealei\** for the presence and distribution of *fmrf*-expressing cells and fiber networks. Late squid embryos and hatchlings were examined by in situ hybridization and immunohistochemistry in whole mounts and tissue sections. All central lobes contained limited numbers of scattered neurons expressing *fmrf*, but the FMRFamide-containing fiber systems were abundant and extensive, mostly present in the neuropil of lobes. Main clusters of neurons were located in the magnocellular and chromatophore lobes of the posterior subesophageal mass (PSM), and in dorsal aspects of the basal lobe (BL). Dense FMRFamide-immunoreactive fibers were particularly seen in the optic lobe (OL), medial and posterior supraesophageal masses (MSM and SPM) often with a commissural organization. The data show that the central lobes of *Doryteuthis pealei* hatchlings have a matured FMRFamide system organized in a limited number of centers, but with widely distributed efferents. This suggests that FMRFamide neurons are already functionally engaged in the late embryo. The localization indicates that control of chromatophores and fin movement are amongst these functions.

## Introduction

FRMFamide belongs to the most abundant regulatory peptides throughout the animal kingdom with the exception of mammals. FMRFamide immunoreactivity has been detected in the oldest branches with complex nervous systems, the Ctenaphores (Moroz et al., 2014), and widely occurs in invertebrates (Fukusumi et al., 2006; López-Vera et al., 2008; Santama and Benjamin, 2000; Sweedler et al., 2000; Walker et al., 2009). This regulatory peptide serves nerve cells as neuropeptide and peripheral endocrine organs as hormone. It has been shown that FMRFamide can modulate neurotransmission between neurons, mediates neural control of peripheral organs, like heart, kidney and intestine, and influences behavior (Bechtold and Luckman, 2007; Bulloch et al., 1988; Di Cosmo and Di Cristo, 1998; Di Cristo et al., 2003; Lehman and Greenberg, 1987). These data indicate that organisms employ FMRFamide for diverse functions in multiple physiological systems.

FMRFamide was first identified in the bivalve mollusk *Macracallista nimbosa* and later in several other molluskan species including cephalopods (Bulloch et al., 1988; di Cosmo and di Cristo, 2006; Price and Greenberg, 1977; Santama and Benjamin, 2000; Sweedler et al., 2000). We recently identified the *fmrf* gene transcript and its peptide products, including FMRFamide, in the North Atlantic long-finned squid *Doryteuthis pealei (*previousy *Loligo pealei)*, and found that the single *fmrf* gene in this cephalopod is regulated by differential expression (Burbach et al., 2014). The level of mRNA transcribed from the *fmrf* gene appeared high in the peripheral stellate ganglion of the giant fiber system, while the FMRFamide peptide was hardly produced. We also noted that the central nervous system of the *Doryteuthis pealei* contained numerous FMRFamide-containing neurons, already before hatching. Embryonic expression has also been observed in the pygmy squid *Idiosepius notoidus* (Wollesen et al., 2010). This suggests that FMRFamide is operating already embryonically and may play a role in the dramatic changes in nervous functions associated with hatching. In this study we therefore aimed to identify the brain systems that use FMRFamide as neurotransmitter around hatching.

## Materials and Methods

### Animals

Live North Atlantic Long-finned squid (*Doryteuthis pealei)* were obtained through the Marine Resources Center of the Marine Biological Laboratory, Woods Hole, MA, USA, and handled according to the ethics guidelines. Late embryos and hatchlings (stages 29 and 30) respectively were obtained from squid egg fingers maintained in tanks with fresh running seawater at 20°C and staged according to Arnold (Arnold, 1965). Embryos (stage 26-29) and hatchlings (stage 30) were harvested in seawater, fixed in 70% ethanol and stored at −20°C for *in situ* hybridization, fixed in Bouins fixative or in 4% paraformaldehyde (PFA) for paraffin embedding and immunohistochemistry.

### In situ hybridization

Ethanol-stored embryos and hatchling squid were embedded in TissueTek and cryosectioned. *In situ* hybridization on 20 μm sections using DIG-labeled RNA probes was performed as described (Smidt et al., 2004). The probe used was a full-length *fmrf* cDNA of *Doryteuthis pealei* (Burbach et al., 2014)

### Immunohistochemistry on sections

Bouin’s-fixed or PFA-fixed embryos and hatchlings were embedded in paraffin and 5 μm sagittal, horizontal, and coronal sections were used for immunocytochemistry as previously described (Grant et al., 1995). Rabbit anti-FMRFamide polyclonal antisera from Chemicon International (AB1917) and from ABCAM (ab10352) were used as primary antibodies. Both antisera were titered for optimal signal-to-background ratio and used at a final concentration of 1:750. If not specified otherwise, antiserum AB1917 (Chemicon International) was used.

Immunoreactivity was detected by DAB-staining using the Vectastain Universal Elite ABC Kit (Pk-6200, Vector Laboratories, Burlingame, CA, USA). As controls for specificity, sections were treated with primary antisera pre-incubated with FMRFamide. Sections were counterstained in 0.2% methyl-green. In one experiment, hatchling sections were treated with a rabbit-derived polyclonal antibody to squid phospho-heavy neurofilament (NFH) (1:500). Separate sections were treated with hematoxyline and eosin (H&E) stain. Photomicrographs were taken with a Zeiss Axiomat microscope system and resulting images adjusted for white balance, brightness, contrast and color level distribution using Photoshop (Adobe) or the Gimp open source software packages.

### Whole mount immunohistochemistry

PFA-fixed squidlets were washed for a day in PBS (containing up to 2% Triton-X 100) then placed in 10% normal goat serum (Invitrogen) for another 24 hours. Rabbit anti-FMRF was added at dilutions of about 1:200 and 1:400 and whole animals were incubated for several days at 10 °C. After rinsing again in PBS, an Alexa-405 anti-rabbit secondary was added and the preparations were shaken overnight at 10 °C. They were rinsed several times in PBS and then mounted directly or in Conray-30 (Mallincrodt) used as an aqueous clearing medium). Animals were imaged as whole mounts in 2D and 3D using a Zeiss LSM780 confocal microscope using its 405, 488 and 633 laser lines, and 10x to 25x objectives.

### Silver staining

Bouin-fixed, paraffin embedded serial sections (10μm) of hatchlings were stained as described by Gregory (Gregory, 1980)Sections were impregnated in 2% protargol solution (pH 2.0) containing copper (1.3gm/65ml) for 12 hours, washed, then developed in 1% hydroquinone containing 2.5% sodium sulfite, washed, gold intensified, dehydrated, cleared and mounted in Permount.

## Results

### Body distribution of FMRFamide immunoreactivity

Confocal microscopic examination of whole-mount preparations of squid hatchlings stained by immunohistochemistry for FMRFamide showed staining in different body parts. Most pronounced staining was in the head region, thorax, heart and vasculature (Fig. 1). These results showed that f*mrf* gene expression had already been activated in peripheral organs and in the nervous system at hatching. Next, we focused on the brain to determine neural sites of gene and peptide expression.

**Fig 1.**
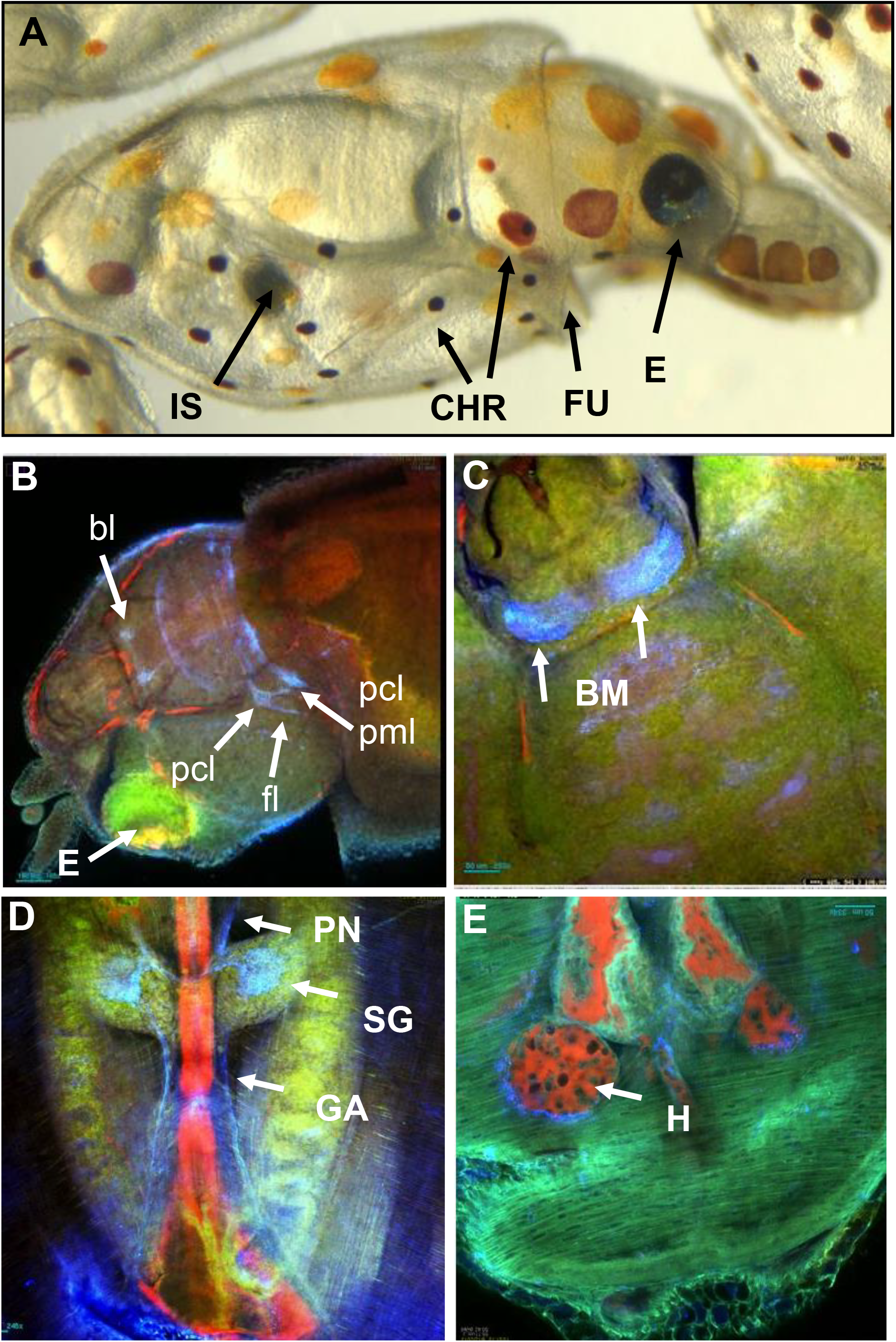
Body distribution of FMRFamide immunoreactivity in a squid hatchling by whole mount immunofluorescence. (A) Overview on the squid hatchling showing eye (E), chromatophores (CHR) and ink sac (IS). (B) FMRFamide-immunoreactive structures in the head region; side view. (C) FMRFamide-immunoreactive structures in the head region; top view. (D) FMRFamide-immunoreactivity in the dorsal body region showing staining of the stellate ganglion (SG), giant axon (GA) and pallial nerve (PN). (E) FMRFamide around the heart (H). Anatomical definitions were according to Young (Young, 1974; Young, 1976; Young, 1977; Young, 1979a; Young, 1979b), Shigeno (Koizumi et al., 2016; Shigeno et al., 2001; Yamamoto et al., 2003). Labeled structures are BM=buccal mass, CHR=chromatophores, E=eye, FU= funnel, GA= giant axon, H=heart, IS=ink sac, PN=pallial nerve, SG=stellate ganglion. Labeled lobes are bl=buccal lobe, fl=fin lobe, pcl=posterior chromatophore lobe. Colors: blue: FMRFamide; red: blood; green: autofluorescence.

### Expression of the *fmrf* gene in brain of late embryos and hatchlings

The CNS complex of the squid consists of distinct ‘masses’, also addressed here as ganglia, containing lobes and interconnected by fiber systems. These ganglia, seen in a silver stained sagittal section of a stage 30 hatchling are organized in the head region (Fig. 2A) and have been anatomically described in great detail for *Doryteuthis pealei* by Young (Young, 1974; 1976; 1976; 1979). We used definitions of anatomical regions according to Young (Young, 1974; Young, 1976; Young, 1977; Young, 1979a; Young, 1979b), Shigeno (Koizumi et al., 2016; Shigeno et al., 2001; Yamamoto et al., 2003).

**Fig 2.**
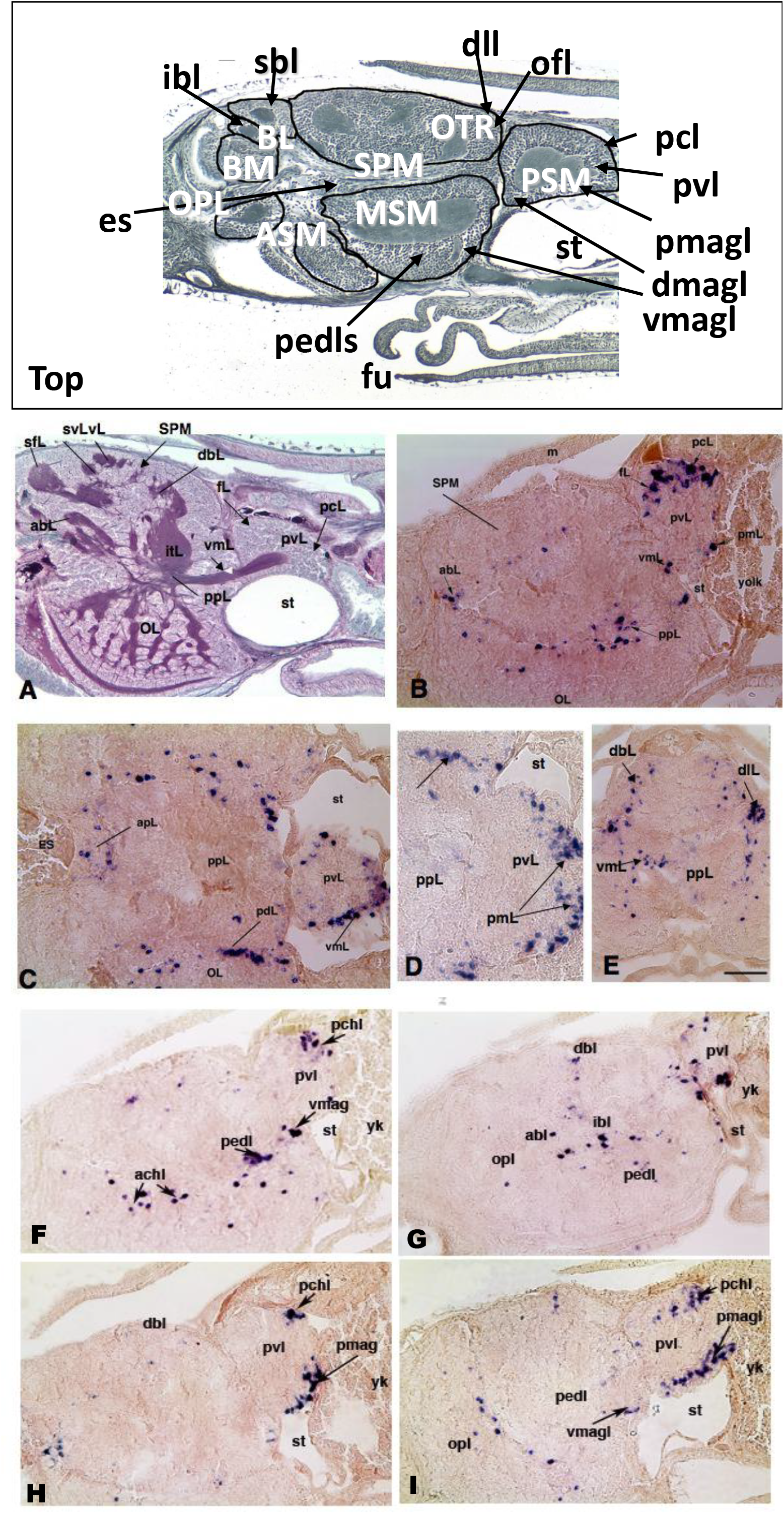
In situ hybridization of *fmrf* gene transcripts in a squid hatchling. (Top) Organization of the central nervous system of the squid embryo depicted by a silver-stained mid-sagittal section of a stage 30 hatchling. The masses shown (in white) are ASM=anterior subesophageal mass; BL=buccal lobes; BM= buccal mass; MSM=medial subesophageal mass OPL=optic lobe; OTR= optic tract region; PSM=posterior subesophageal mass; SPM=supraesophageal mass. (A) For further detail, a silver stained sagittal section of a stage 30 hatchling zoomed in on the central nervous system of the squid embryo. (B-I) In situ hybridization of cryostat sections of a stage 30 embryo showing expression of *fmrf* mRNA in cells in different brain regions. (B, F-I) Saggital sections through the mid-region showing scattered and clustered cells expressing *fmrf* mRNA. (C-D) Horizontal sections at different magnifications to show clusters of *fmrf* expressing cells in the PSM. (E) Coronal section to show *fmrf* expression in the SPM. Abbreviations are: abL= anterior basal lobe; apL=anterior pedal lobe; dbL=dorsal basal lobe; dlL=dorsal lateral lobe; ES=esophagus; fL fin lobe; ibL=inter basal lobe; m=mantle; OL=optic lobe; pcL=posterior chromatophore lobe; pdL=peduncle lobe; pedl=pedal lobe; pmagl=posterior magnocellular lobe; ppL=posterior pedal lobe; pvL=palliovisceral lobe; sfL=superior frontal lobe; st=statocyst; svL= subvertical lobe; vmagl=ventral magnocellular lobe; vL=vertical lobe; yk=yolk. Scale bar in panel E: 50u abl=anterior basal lobe; achl= anterior chromatophore lobe; dbl=dorsal basal

We examined the distribution of *fmrf-*expressing neurons in the developing CNS by *in situ* hybridization (Fig. 2). No expression of the gene was detected in embryos of stage 26 and before. Stage 29 embryos and stage 30 hatchlings displayed distinct patterns of expression that were detailed further on sagittal, horizontal, and coronal sections of squid with no significant difference between these stages (Fig S1, Supplement).

Overviews obtained by serial sections showed that the expression of the *fmrf* gene displayed a pattern of sparse and scattered neurons over all brain masses with a few clusters of neurons (Fig. 2). Three areas stood out in density of *fmrf*-expressing neurons. Firstly, the largest clusters of *fmrf*-expressing neurons were found in the posterior subesophageal mass (PSM) containing the palliovisceral lobe (pvl). This lobe contained several groups of neurons overlapping the dorsal aspect of the posterior chromatophore lobes (pcL, Fig. 2B, S1), fin lobes (fL, Fig. 2B, S1) and the ventral complex of magnocellular lobes (vmagl, Fig. 2B, C, E) as well as the posterior magnocellular lobes (pmagl, Fig. 2B, D).

Secondly, very dense staining *fmrf*-expressing neurons were detected in the dorsal basal and dorsal lateral lobes of the basal lobe (BL) complex (dbL, dlL Fig. 2E). Anterior neural structures were sparse in *fmrf*-expressing neurons. Thirdly, the optic lobe (OL) contained scattered *fmrf*-expressing neurons in the medulla and a post-pedal lobe (ppL Fig. 2B). Notably, *fmrf* gene expression was present in neurons intermediate to the OL and the central brain masses, visible in serial coronal sections (Fig S2)

The data indicate that the *fmrf* gene is transcriptionally active in limited and scattered sets of distinct neurons. Neurons are distributed over most brain lobes with some regions particularly rich in *fmrf*-expressing neurons such as the palliovisceral, magnocellular, and basal lobes of the PSM. Next, we determined the presence and distribution of the FMRFamide tetrapeptide.

### FMRFamide in ganglia, lobes and fiber tracts

Immunohistochemistry on sections of hatchlings and whole-mount preparations revealed brain structures containing FMRFamide peptide (Fig. 3 and 4). Whole mounts analyzed by immunofluorescence and confocal microscopy displayed intense staining in the head region confined to lobes and fiber-like structures (Fig. 3, Fig. S1). At the level of the PSM, dense immunoreactivity in neurons and fiber networks were located in the posterior region containing the chromatophore lobe, fin lobe and magnocellular lobe (Fig 3C, D). This corresponded to the presence of *fmrf* mRNA in these lobes (Fig. 2). In addition, FMRFamide immunoreactivity appeared in dense fiber tracts associated with these lobes that displayed commissural connectivity (Fig 3C, D). Fine fiber networks were observed in the optic lobe (Fig 3D). These structures were further examined by immunohistochemistry on serial sections.

**Fig 3.**
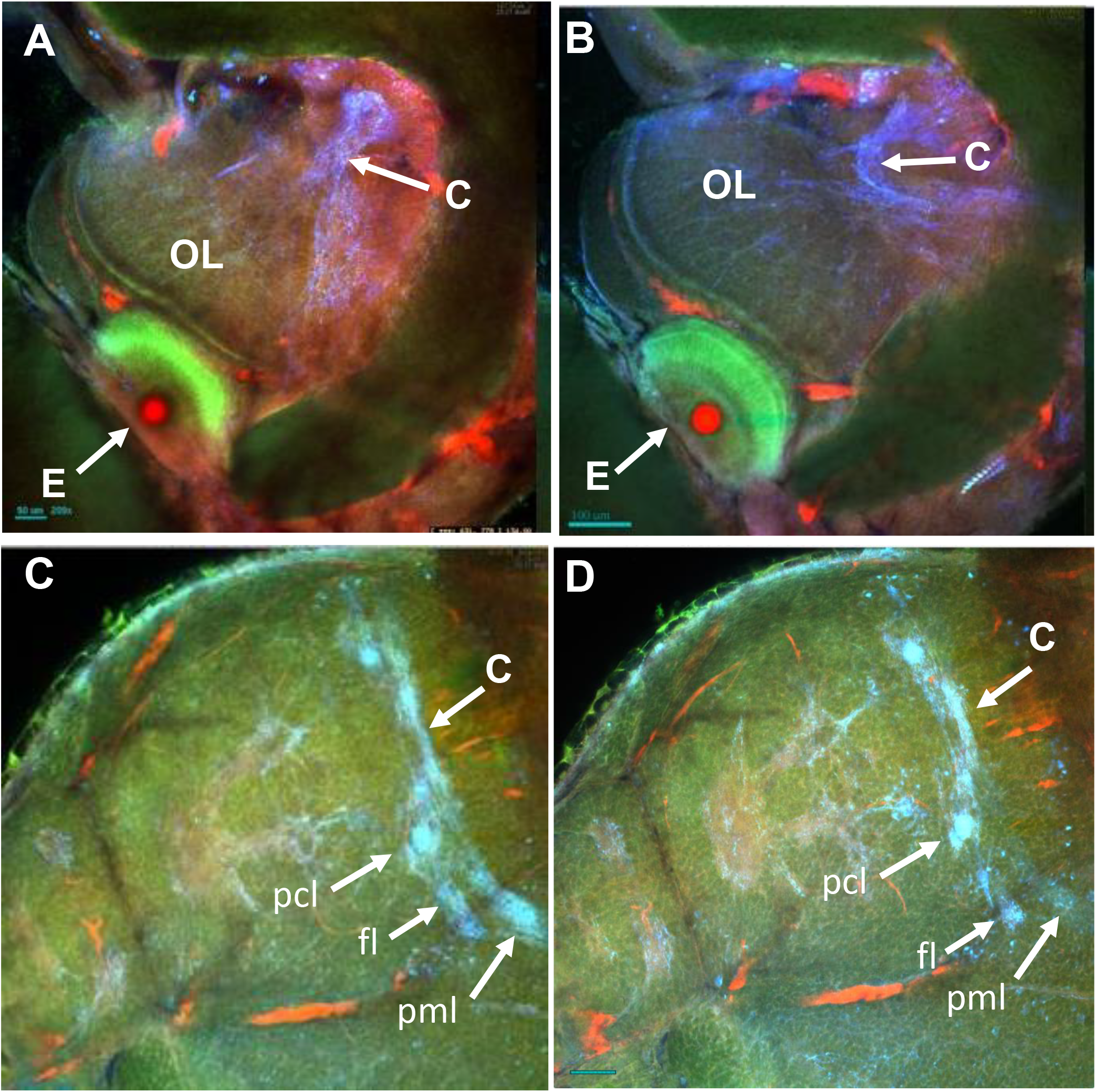
Immuofluorescence of FMRFamide-immunoreactivity in the head region of hatchling analyzed by confocal microscopy. (A,B) Angular view on the head showing FMRFamide immunofluorescent fibers (blue) in the optical lobe (OL) neighboring the eye (E) and commissural connectivity (C) in the region. (C,D) View on the chromatophore lobes showing dense labeling in the posterior chromatophore lobe (pcl), fin lobe (fl) and posterior magnocellular lobe (pml). Colors: blue: FMRFamide; red: blood; green: autofluorescence. More detail is shown in a movie reconstruction of confocal analysis through the head region in the Supplementary data (Fig. S1).

**Fig 4.**
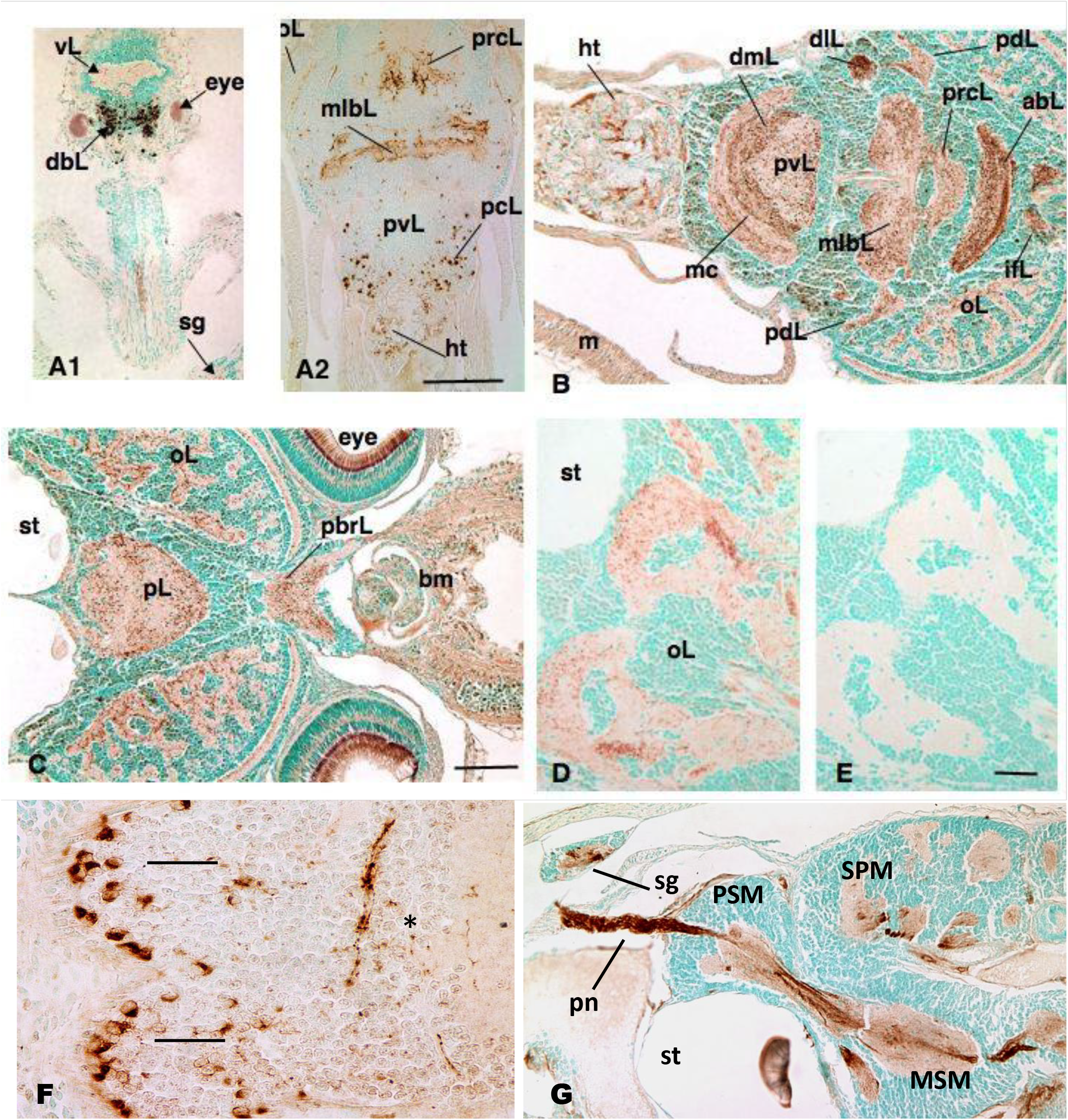
FMRFamide immunostaining on the central nervous system of squid hatchlings. A-C: Paraformaldehyde-fixed paraffin sections (except A2, Bouins fixed) from dorsal to ventral at same magnification immunostained with FMRFamide antibody. D-E: Paraffin sections at similar level as C at higher magnification to show FMRFamide expression in medulla of optic lobe (D) compared with an adjacent section pre-absorbed with FMRF peptide (E) demonstrating specificity of FMRFamide staining. F-G: horizontal section through the PSM displaying FMRFamide-positive neurons of the magnocellular lobe (F) and a sagital section showing the pallial nerve tract leaving the PSM (G). Legend to abbreviations: same as in Fig. 2. including additional: bm= buccal mass; dmL = dorsal magnocellular lobe; ht= heart: ifL = inferior frontal lobe; mlbL = mediolateral basal lobe complex; mc= magnocellular commissure; pal = pallial nerve; pL = pedal lobe; pbrL= pre-commissural lobe; pn=pallial nerve; sg=stellate ganglion; st=statocyst. Scale bars: A1,A2: 100u; B,C: 50u; D,E: 20u

In the most dorsal horizontal section through the head region, which also included a portion of the SG (Fig. 4A1), dense clusters of neurons in the dorsal basal lobes exhibited the most intense FMRFamide immunoreactivity, consistent with expression of the *fmrf* mRNA seen in Fig. 2 and Supplemental Fig S2 and the whole mount preparation (Fig. 3 and Fig. S1).

In more ventral sections also including the PSM (Fig. 4A2), neuronal cell labeling was most intense in the dorsal regions of the palliovisceral lobe, i.e., the fin lobes and posterior chromatophore lobes. In this section and in Fig. 4B the cells and nerve fibers in the region caudal to the palliovisceral lobe, were also immunoreactive. The intraganglionic spaces of the medio/lateral basal lobes (mlbL) and precommissural lobe (prc) were filled with FMRFamide-labeled fibers, tracts and neuropil. The fiber tracks corresponded to those visible in immunostained whole mount preparations (Fig 3).

In contrast to the PSM, the SPM contained few FMRFamide-immunoreactive neurons, but displayed immunoreactivity principally in intraganglionic fibers and tracts of several SPM lobes such as the optic, frontal, vertical and brachial lobes (Fig. 5). These regions are involved in memory and learning of tactile and visual information (Williamson and Chrachri, 2004). FMRFamide-immunoreactive fibers were dense in the neuropil of the SPM, particularly in the optic tract region that interphases the optic lobes and central brain mass.

**Fig 5.**
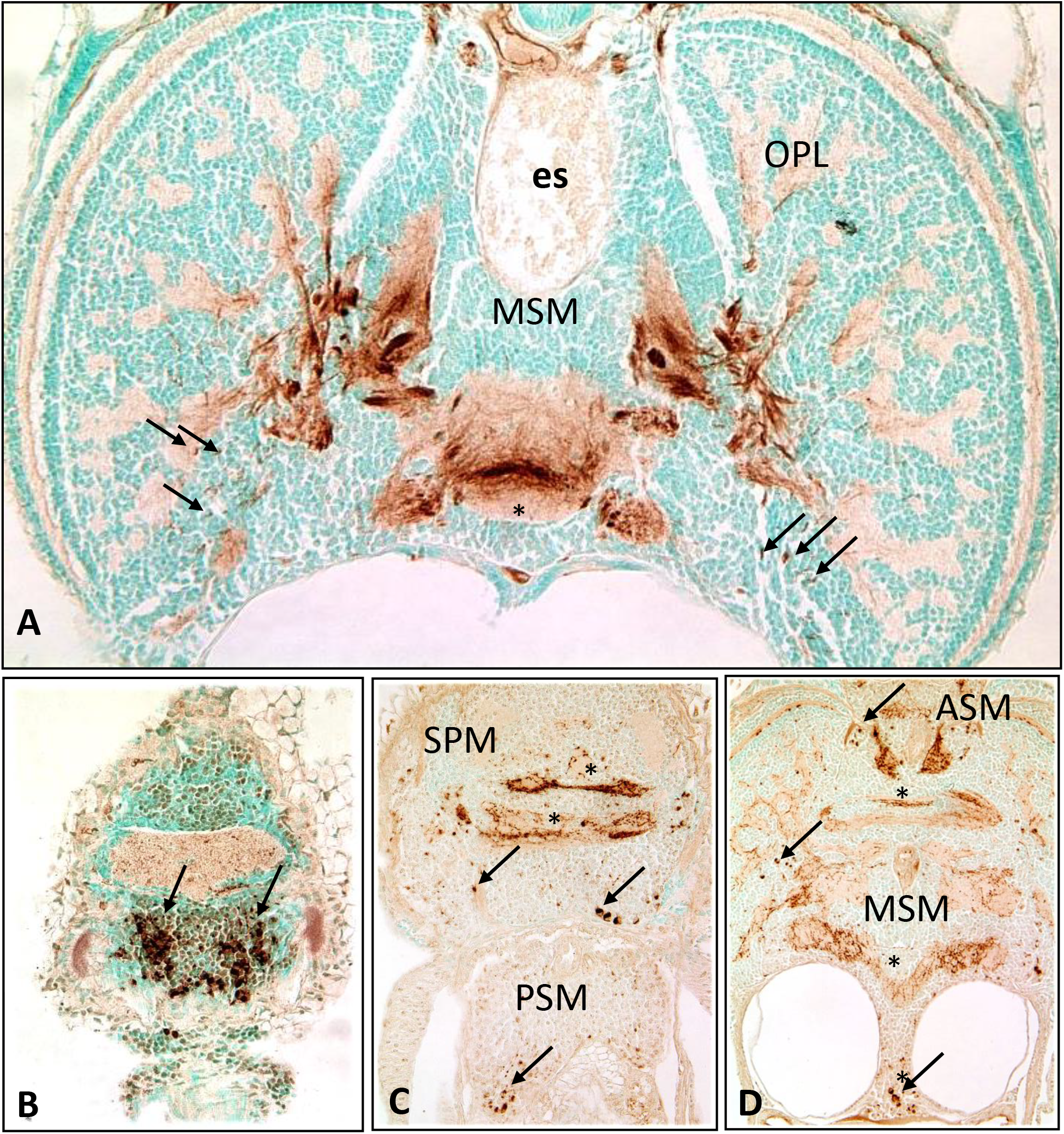
FMRFamide immunoreactive tracts of the nervous system of the squid hatchling. (A) Coronal section through the optic lobe region showing dense fiber systems connected to the optic lobe (OL) and the medial subesophageal mass (MSM). Furthermore, dense fiber networks are present in the MSM neuropil. Arrows indicate FMRFamide-positive cell bodies. (B-D) Horizontal sections showing the organization of FMRFamide-immunoreactive tracts at different levels. Note the commissural structures in the SPM, MSM and anterior subesophageal mass (ASM), labeled *. Arrows indicate FMRFamide immunoreactive cell bodies.

The middle subesophageal mass (MSM) contained areas with several *fmrf*-expressing neurons and FMRFamide immunoreactive fibers and neuropils. One area was at the position of the pedal lobes (Fig. 5 and Fig. 2C). Lateral areas of the MSM bordering the optic lobes contained *Fmrf-expressing* neurons. Dense fiber networks and commissures in these areas and in the optic tract suggested that these neurons were connected to the optic lobes (Fig. 5C, Fig. 2). The most medial structures of the optic lobes contained neurons that expressed the *fmrf* gene (Fig. S1). Lateral areas were devoid of *fmrf* expression. Interestingly, a fine fiber tract was observed in the outer neuropil layer surrounding the optic lobes while the medulla was filled with a rich supply of FMRFamide fiber immunoreactivity (Fig. 5).

## Discussion

This study shows that the *fmrf* gene is expressed widely in the central nervous system of the squid *Doryteuthis pealei* around hatching and leads to biosynthesis of FMRFamide, the biologically active gene product. This contrast *fmrf* gene expression in the stellate ganglion, the major peripheral ganglion that mediates mantle contraction, which contains abundant *fmrf* mRNA, but is very low in FMRFamide peptide at this stage (Burbach et al., 2014). The neural expression patterns indicate that the biological functions exerted by FMRFamide in the central nervous system are already operating during hatching, suggesting that they may be required for the physiological control of the squid hatchling.

The most notable characteristic of brain expression of fmrf is the restricted expression of the gene to limited and defined neurons, but the very abundant presence of FMRFamide-immunoreactive fibers throughout the central brain masses. This suggests that the neurons expressing the *fmrf* gene are far-reaching and may serve important functions in integrating various central systems. Particularly, the large central neuropil of ASM, MSM and SPM were filled with dense FMRFamide immunoreactivity and commissural-like structures were visible (Fig. 1, Fig. 4 and Fig. 5). This is further illustrated by the optic lobe. While there were few neuronal cell bodies expressing FMRFamide in the optic lobes (Fig. 3 and Fig 5), these lobes contained dense FMRFamide-immunoreactive fiber networks in the medulla and cortical plexiform layer as well as the optic tract fibers (Fig 3 and Fig 5A). This suggests that the optic lobes may receive FMRFamide-containing afferents from other lobes.

The overall distribution of FMRFamide in Doryteuthis pealei corresponds broadly with the distribution of FMRFamide-positive cells found in the nervous systems of other cephalopods, *Sepia officinalis* and *Idiosepius notoides*, as far as comparable in the published data (Aroua et al., 2011; Wollesen et al., 2008; Wollesen et al., 2010). In particular, the central palliovisceral lobe, chromatophore and magnocellular lobes seem to be shared as lobes with relatively dense FMRFamide neurons. This suggests that the engagement of FMRFamide in neurobiologial functions associated with these neurons is conserved.

Several clusters of neurons and associated fiber systems could be related to peripheral functions and suggest the involvement of FMRFamide in these functions. Firstly, the chromatophore lobes exhibited the most robust expression of the *fmrf* gene and FMRFamide peptide. Chromatophores are controlled before hatching (Messenger, 2001). This suggests that FMRFamide may be a main player in the regulation of chromatophores which role is already fully established before hatching. Our data are consistent with several studies on *Sepia officinalis* highlighting the role of FMRFamide peptides in the regulation of color changes during development and in the adult (Aroua et al., 2011; Loi and Tublitz, 2000; Loi et al., 1996). Notably, FMRFamide immunoreactivity is present in all trajectories of the neural pathways controlling chromatophore function (Aroua et al., 2011). FMRFamide modulates and supplements glutamatergic control over the fast and slow transformations of color patterns in this species (Loi and Tublitz, 2000; Loi et al., 1996). This FMRFamide system may belong to one of the first neural networks in the squid brain that are activated early during embryonic development (Williamson and Chrachri, 2004).

Secondly, the fin lobes contained dense clusters of *fmrf* gene and peptide expression. We also observed FMRFamide immunoreactivity in the mantle at the level of the fins (not shown). Fibers reach the fins through the fin nerve that is formed by descending fibers in the pallial nerve from the brain, passing the stellate ganglion and running as fine fiber structures in association with the giant axons. FMRFamide reactivity was observed in the pallial nerve (Fig 1C and Fig. 4G), in the neuropil of the stellate ganglion (Fig 4G) and in association with the giant axon (Burbach et al., 2014). This anatomical organization supports the notion that FMRFamide plays a role in the fine control of fin movement (Williamson and Chrachri, 2004).

This study shows that the central ganglia contain FMRFamide in multiple brain systems underpinning the concept that FMRFamide is widely engaged in physiological systems of invertebrates. It illustrates the pleiotropy displayed by the *fmrf* gene. In hatchlings of *Dorytheuthis pealei* amongst these systems are those that contribute to control of chromatophores and fin movement, functions that are fully operative in the late embryo just before hatching and required after hatching.

## Authors contributions

JPHB conceived of the study, designed the study, coordinated the study, carried out parts of the lab work, and drafted the manuscript; PG participated in the design of the study, carried out parts of the lab work and helped to draft the manuscript; SS and LK carried out the immunofluorescence confocal analyses; AJCGMH carried out parts of the lab work; HCP participated in the design of the study and helped to draft the manuscript. All authors gave final approval for publication.

## Ethics

Animals were collected, maintained, handled and treated according to the ethics guidelines of the Marine Resource Center, Marine Biological Laboratory (MBL), Woods Hole, MA, USA.

---

**Fig. S1:**
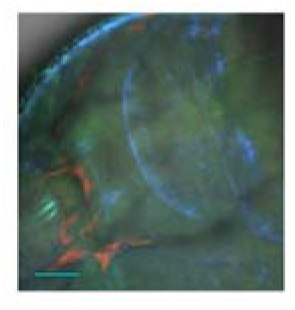
Immunofluorescence of FMRFamide-immunoreactivity in the head region of a squid hatchling presented as a movie reconstruction of serial confocal microscopy pictures.

**Fig S2:**
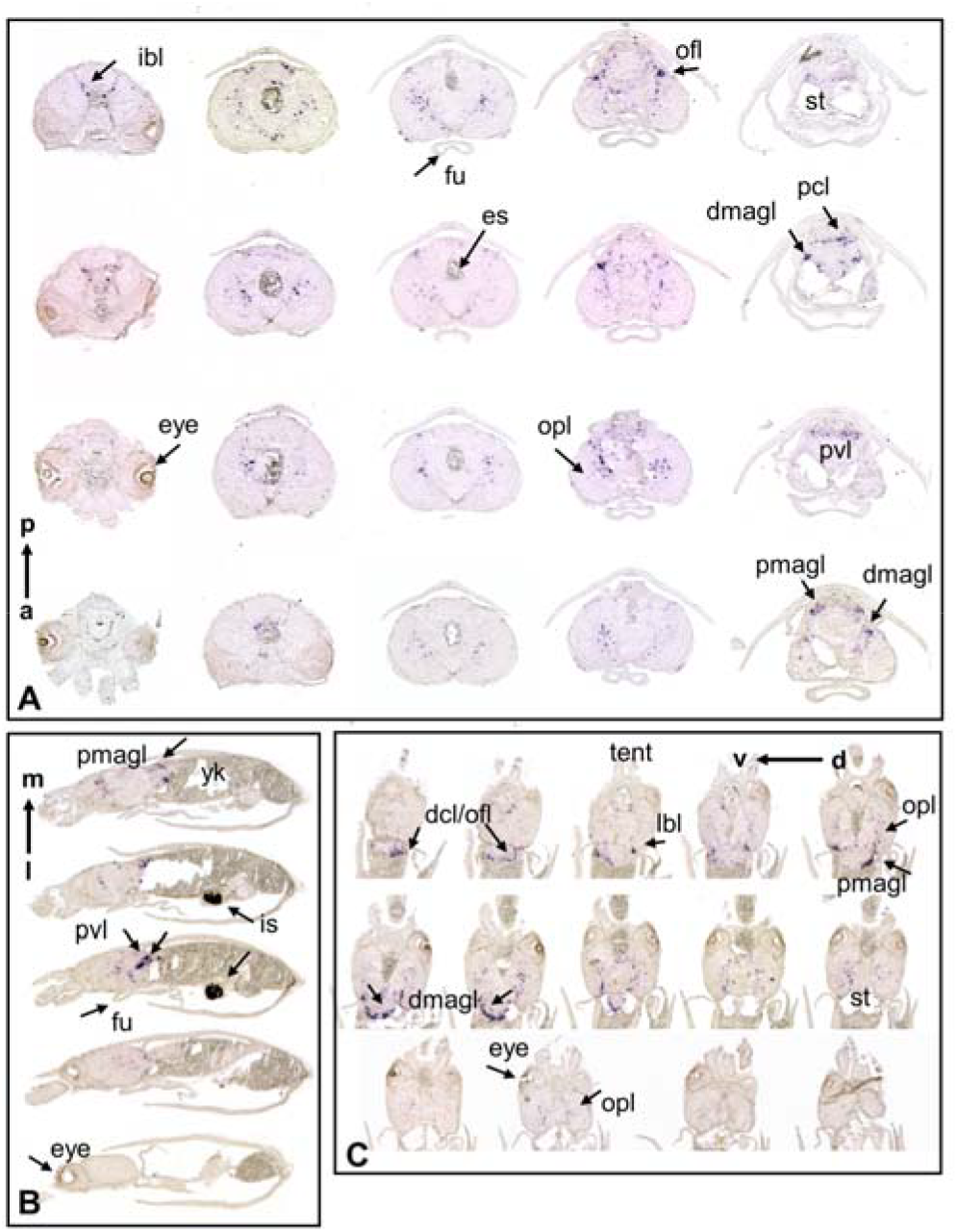
In situ hybridization of *fmrf* mRNA in coronal (A), sagittal (B) and horizontal (C) sections of stage 29 squid embryos. Abbreviations are: dcl=dorsal chromatophore lobe; dmagl=dorsal magnocellular lobe; es=; eye=eye; fu=funnel; ibl=; ofl=olfactory lobe; opl=optic lobe; pcl=posterior chromatophore lobe; pmagl=posterior magnocellular lobe; pvl= palliovisceral lobe; st= statocyst; tent=tentacles;

