## Supplementary figures and images for "*FMRF* Gene Expression in the Nervous System of the Squid *Doryteuthis Pealei*\* Hatchling"

### Supplemental figure S2

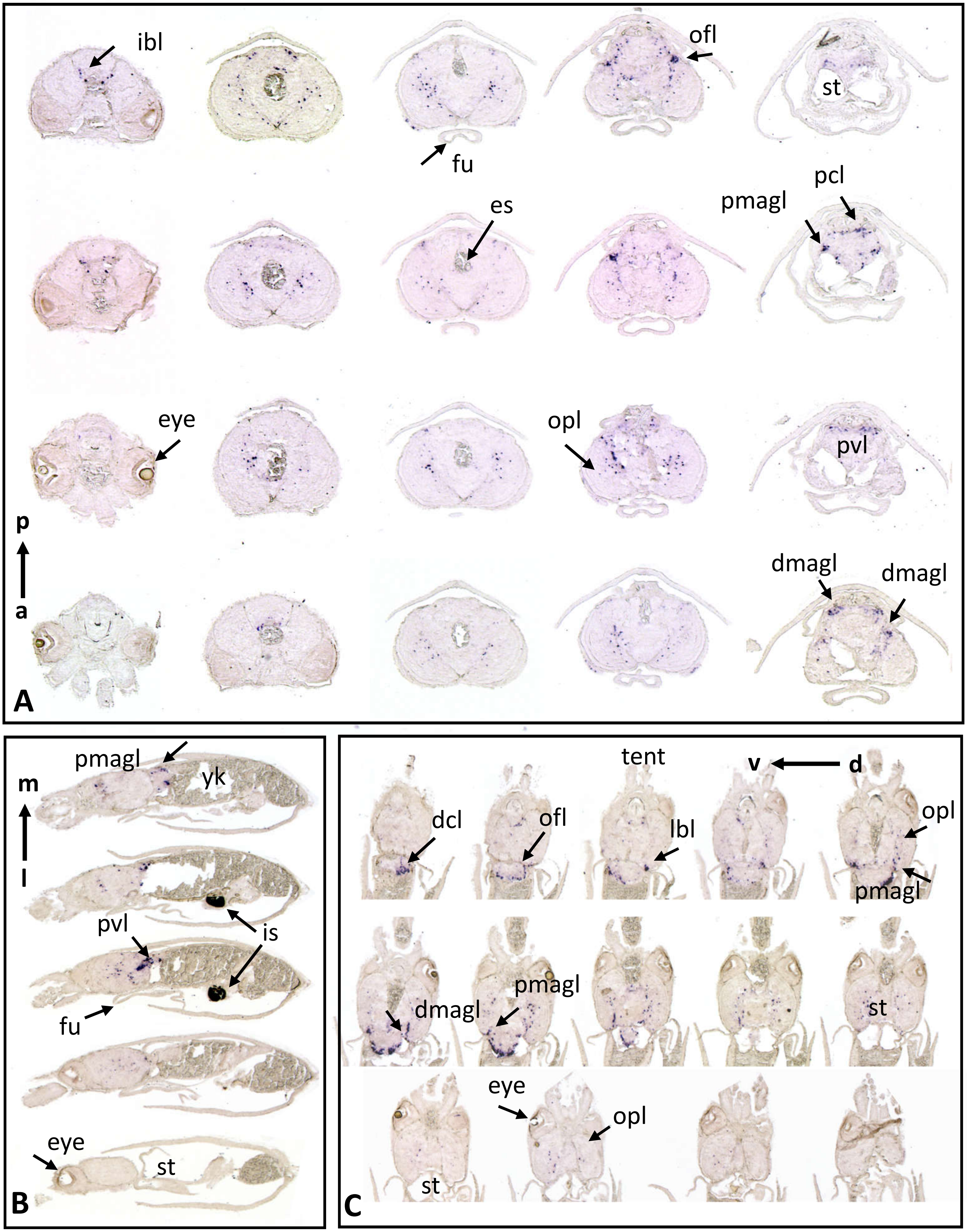


Supplemental figure S2
